# A two-component protein condensate of EGFR and Grb2 regulates Ras activation at the membrane

**DOI:** 10.1101/2021.12.12.472247

**Authors:** Chun-Wei Lin, Laura M. Nocka, Brittany Stinger, Joey DeGrandchamp, Nugent Lew, Steven Alvarez, Henry Phan, Yasushi Kondo, John Kuriyan, Jay T. Groves

## Abstract

We reconstitute a phosphotyrosine-mediated protein condensation phase transition of the ∼200 residue cytoplasmic tail of the epidermal growth factor receptor (EGFR) and the adaptor protein, Grb2, on a membrane surface. The phase transition depends on phosphorylation of the EGFR tail, which recruits Grb2, and the dimerization of Grb2, which provides the crosslinking element for condensation with EGFR. The Grb2 Y160 residue plays a structurally critical role in dimer formation, and phosphorylation or mutation of Y160 prevents EGFR:Grb2 condensation. By extending the reconstitution experiment to include the guanine nucleotide exchange factor, SOS, and its substrate Ras, we further find that EGFR condensation controls the ability of SOS to activate Ras. These results identify an EGFR:Grb2 protein condensation phase transition as a regulator of signal propagation from EGFR to the MAPK pathway.

## Introduction

Recently, a class of phenomena known as protein condensation phase transitions have begun to emerge in biology. Originally identified in the context of nuclear organization (1) and gene expression (2), a distinct two-dimensional protein condensation on the cell membrane has now been discovered in the T cell receptor (TCR) signaling system involving the scaffold protein LAT (3–5). TCR activation results in phosphorylation of LAT on at least four distinct tyrosine sites, which subsequently recruit the adaptor protein Grb2 and the signaling molecule PLCγ via selective binding interactions with their SH2 domains. Additional scaffold and signaling molecules, including SOS, GADS, and SLP76 are recruited to Grb2 and PLCγ through further specific protein-protein interactions (6, 7). Multivalency among some of these binding interactions can crosslink LAT molecules in a two-dimensional bond percolation network on the membrane surface. The resulting LAT protein condensate resembles the nephrin:NCK:N-WASP condensate (8) in that both form on the membrane surface under control of tyrosine phosphorylation and exert at least one aspect of functional control over signaling output via a distinct type of kinetic regulatory mechanism (9–11). The basic molecular features controlling the LAT and nephrin protein condensates are common among biological signaling machinery and other similar condensates continue to be discovered (12, 13). The LAT condensation shares downstream signaling molecules with the EGF receptor (EGFR) signaling system, raising the question if EGFR may participate in a signaling-mediated protein condensation itself.

EGFR signals to the mitogen-activated protein kinase (MAPK) pathway and controls key cellular functions including growth and proliferation (14–16). EGFR is a paradigmatic model system in studies of signal transduction and immense collective scientific effort has revealed the inner workings of its signaling mechanism down to the atomic level(17). EGFR is autoinhibited in its monomeric form. Ligand-driven activation is achieved through formation of an asymmetric receptor dimer in which one kinase activates the other to phosphorylate the nine tyrosine sites in the C-terminal tails (17, 18). There is an obvious conceptual connection between EGFR and the LAT signaling system in T cells. The ∼200 residue long cytoplasmic tail of EGFR resembles LAT in that both are intrinsically-disordered and contain multiple sites of tyrosine phosphorylation that recruit adaptor proteins upon receptor activation (19). Phosphorylation at tyrosine residues Y1068, Y1086, Y1148, and Y1173 in the EGFR tail creates sites to which Grb2 can bind via its SH2 domain. EGFR-associated Grb2 subsequently recruits SOS, through binding of its SH3 domains to the proline-rich domain of SOS. Once at the membrane, SOS undergoes a multi-step autoinhibition release process and begins to catalyze nucleotide exchange of RasGDP to RasGTP, activating Ras and the MAPK pathway (20).

While these most basic elements of the EGFR activation mechanism are widely accepted, larger-scale features of the signaling complex remain enigmatic. A number of studies have reported higher-ordered multimers of EGFR during activation (21–24). Structural analyses and point mutations on EGFR have identified a binding interface enabling EGFR asymmetric dimers to associate (24), but the role of these higher-order assemblies remains unclear. At the same time, many functional properties of the signaling system remain unexplained as well. For example, EGFR is a frequently altered oncogene in human cancers, and drugs (including tyrosine kinase inhibitors) targeting EGFR signaling have produced impressive initial patient responses (25). All too often, however, these drugs fail to offer sustained patient benefits, in large part because of poorly understood resistance mechanisms (26). Physical aspects of the cellular microenvironment have been implicated as possible contributors to resistance development (27) and there is a growing realization that EGFR possesses kinase-independent (e.g. signaling independent) pro-survival functions in cancer cells (28). These points fuel speculation that additional layers of regulation over the EGFR signaling mechanism exist, including at the level of the receptor signaling complex itself.

Here we report that EGFR undergoes a protein condensation phase transition upon activation. We reconstitute the cytoplasmic tails of EGFR on supported bilayers and characterize the system behavior upon interaction with Grb2 and SOS using total internal reflection fluorescence (TIRF) imaging. This experimental platform has been highly effective revealing both phase transition characteristics and functional signaling aspects of LAT protein condensates (4, 5, 10, 29–31). Published reports on the LAT system to date have emphasized SOS (or the SOS PR domain) as a critical crosslinking element. Titrating the SOS PR domain into an initially homogeneous mixture of phosphorylated LAT and Grb2 reveals a sharp transition to the condensed phase, which we have also observed with the EGFR:Grb2:SOS system. Under slightly different conditions, however, we report observations of an EGFR:Grb2 condensation phase transition without any SOS or other crosslinking molecule. We show that crosslinking is achieved through a Grb2 dimer interface. Phosphorylation on Grb2 at Y160 as well as a Y160E mutation (both known to disrupt the Grb2 dimer (32, 33)) are observed to prevent formation of EGFR condensates.

The consequence of EGFR condensation on downstream signaling is characterized by mapping the catalytic efficiency of SOS to activate Ras as a function of EGFR condensation state. SOS is the primary Ras guanine nucleotide exchange factor (GEF) responsible for activating Ras in the EGFR to MAPK signaling pathway (34–37). At the membrane, SOS undergoes a multi-step process of autoinhibition release before beginning to activate Ras. Once fully activated, SOS is highly processive and a single SOS molecule can activate hundreds of Ras molecules before disengaging from the membrane (38–40). Autoinhibition release in SOS is a slow process, which necessitates that SOS be retained at the membrane for an extended time in order for Ras activation to begin (5, 10). This delay between initial recruitment of SOS and subsequent initiation of its Ras GEF activity provides a kinetic proofreading mechanism that essentially requires SOS to achieve multivalent engagement with the membrane (e.g. through multiple Grb2 or other interactions) in order for it to activate any Ras molecules.

Experimental results described here reveal that Ras activation by SOS is strongly enhanced by EGFR condensation. Calibrated measurements of both SOS recruitment and Ras activation confirm enhanced SOS catalytic activity on a per-molecule basis, in addition to enhanced recruitment to the condensates. These results suggest that a Grb2-mediated EGFR protein condensation phase transition is a functional element controlling signal propagation from EGFR downstream to the MAPK signaling pathway.

## Results

### Reconstitution of a membrane-surface EGFR:Grb2 protein condensate

We reconstitute an EGFR:Grb2 protein condensation phase transition on supported membranes using the C-terminal region of EGFR (residues 991-1186, where the signal peptide of EGFR is not included in numbering), which is expressed along with a His_6_ tag and a SUMO tag at the N-terminal end of the construct. Throughout the text, we refer to this construct of the cytoplasmic tail of EGFR as the “EGFR^TAIL^”. EGFR^TAIL^ is anchored on the supported lipid bilayer through binding between its His tag and Ni-NTA lipid (incorporated into the supported bilayer at 4% molar ratio with DOPC lipid as the primary component) following previously published methods (5, 10). The SUMO fusion tag is used to increase the expression yield of the disordered EGFR^TAIL^ and acts as the spacer between the surface of the bilayer and the tail of EGFR, where the native kinase domain of EGFR would normally be positioned. The EGFR^TAIL^ is fluorescently labeled with Alexa Fluor 488 through the maleimide chemistry and we achieve 66% labeling efficiency. EGFR^TAIL^ constructs anchored to the supported membrane undergo simple Brownian motion (D= 1.76 µm^2^/s, measured by the fluorescence recovery after photobleaching experiments, Figure S1) and appear uniformly distributed on the membrane surface under total internal reflection fluorescence (TIRF) microscopy. The lateral density of the EGFR^TAIL^ on the membrane can be precisely measured by fluorescence correlation spectroscopy (FCS) (5, 41) and is controlled between 50 to 3000 molecules/µm^2^ in the experiments presented here. To maintain the EGFR^TAIL^ in its active, phosphorylated state, a separate Src-family kinase protein, Hck, is tethered to the membrane (also through His tag chemistry) and ATP is provided in solution. Src kinases do not normally phosphorylate the EGFR^TAIL^ efficiently (19), but in this system the pre-incubation with Hck suffices to ensure robust phosphorylation of the tail. A similar strategy was used previously to maintain LAT in its phosphorylated state for condensation phase transition studies (5, 30, 42). Grb2 and, in later experiments, SOS are added to the pre-phosphorylated EGFR on supported membranes to initiate the condensation phase transition.

Upon addition of Grb2, the EGFR^TAIL^ is observed to undergo a robust condensation phase transition, as illustrated schematically in Figure 1A. Images tracking the phase transition process (Figure 1B) show an initially homogeneous distribution of the EGFR^TAIL^, which undergoes a coarsening process after injection of Grb2 (6 µM) to form a dense phase (bright in these images) interspersed with a sparse phase containing a very low density of monomeric EGFR^TAIL^. The domains exhibit undulating shapes and a sharp discontinuity in EGFR^TAIL^ density at the boundaries of the two phases. The Grb2-driven condensation of EGFR^TAIL^ occurs over a range of initial densities of the tails to form condensed phases of similar density, but different overall area coverage. At lower EGFR^TAIL^ initial density, isolated circular EGFR^TAIL^:Grb2 condensates are observed while fully percolating domain patterns are observed at higher densities (Figure 1C). The presence of both Grb2 and EGFR^TAIL^ in the condensates is confirmed by separately labeling them with Alexa Fluor 488 (EGFR^TAIL^) and Alexa Fluor 647 (Grb2), respectively and performing two-color imaging (Figure 2).

**Figure 1.**
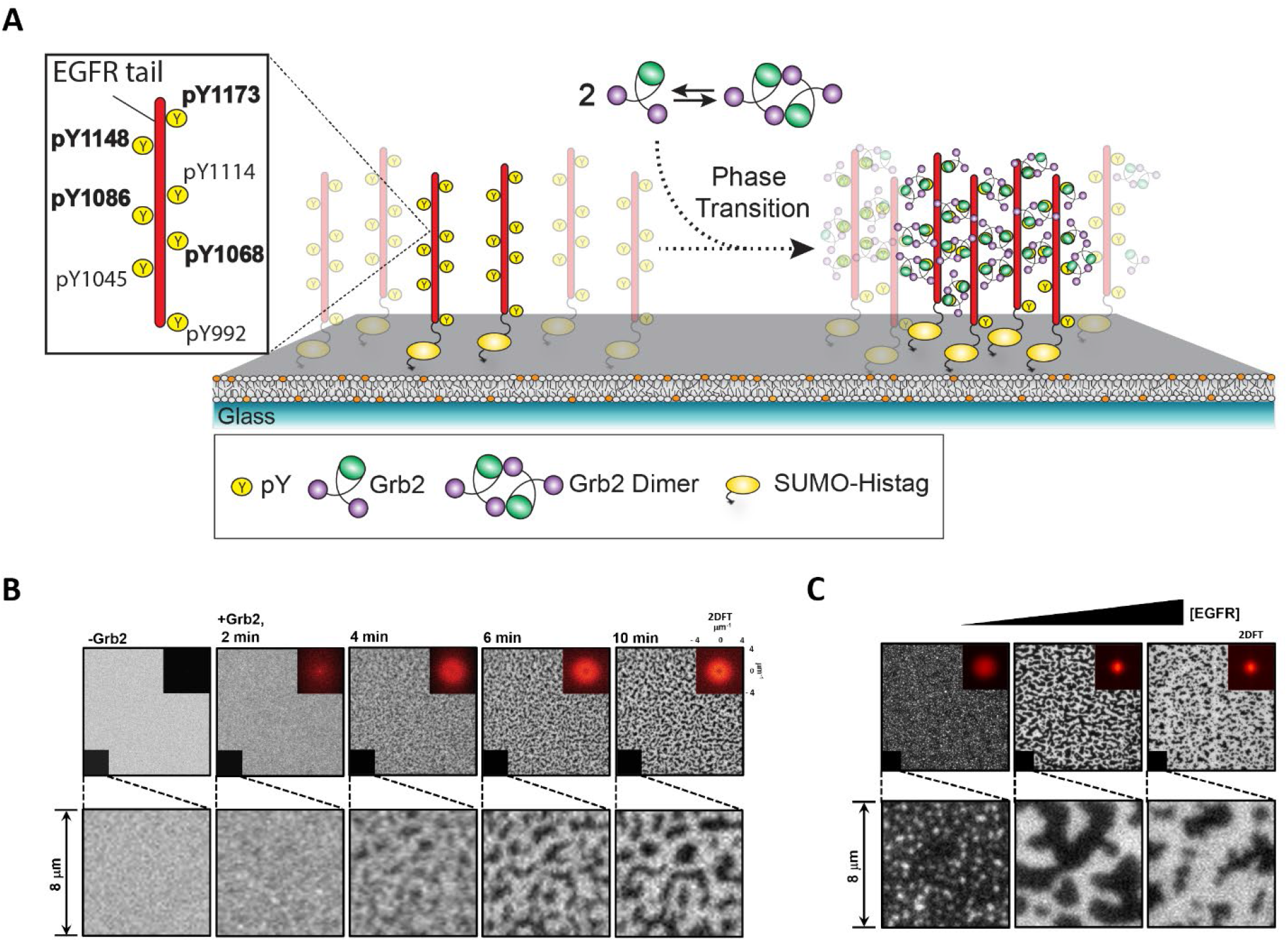
EGFR^TAIL^ phase transition induced by the dimerization of Grb2. (A) The schematic of the reconstitution experiment. The phosphorylated EGFR^TAIL^ is tethered on the supported membrane. EGFR^TAIL^ has several tyrosine sites that can be phosphorylated including Tyr 1068, Tyr 1086, Tyr 1148 and Try 1173 related to Grb2 recruitment. The monomeric Grb2 is shown to be in equilibrium with the Grb2 dimer. Grb2 is injected to the bilayer and recruited by EGFR^TAIL^. The dimerization of Grb2 drives the assembly of EGFR^TAIL^:Grb2 into phase transition. EGFR^TAIL^ is initially in the state where it is mobile on the supported bilayer (left). EGFR^TAIL^ becomes part of the protein condensate after the phase transition is triggered (right). (B) The TIRF images of Alexa Fluor 488 labeled EGFR^TAIL^ on the supported bilayer. EGFR^TAIL^ undergoes phase transition in 2 minutes after the addition of Grb2. The protein condensate of EGFR^TAIL^:Grb2 stabilizes in 10 minutes. The bottom figures are 8-μ m by 8-μ m zoom-in images of the lower-left corner in the upper images. The inserted figures at the upper right corner are the 2D Fourier Transform of the TIRF images. The phase transition shows the contribution at the low spatial frequencies. (C) The size of EGFR^TAIL^:GRB2 condensate is modulated by the density of EGFR^TAIL^ from 50 to 3000 molecules/µm^2^. EGFR^TAIL^:Grb2 condensate varies from small clusters (left) to huge domains (right).

**Figure 2.**
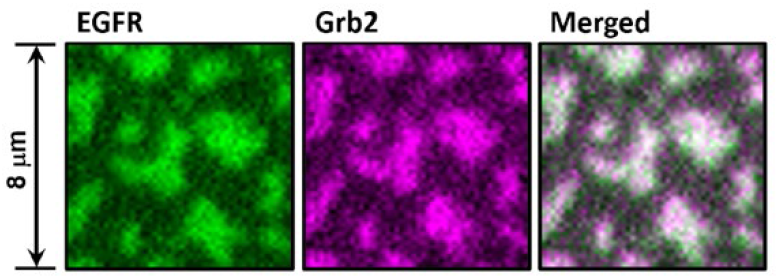
The colocalization between EGFR^TAIL^ and Grb2 after EGFR^TAIL^:Grb2 phase transition is achieved. Unlabeled Grb2 mixed with Alexa Fluor 647 labeled Grb2 (0.2 μM, labeling efficiency of 63%) was added to phosphorylated Alexa Fluor 488 labeled EGFR^TAIL^ to induce the phase transition of EGFR^TAIL^:Grb2. The TIRF images were taken 10 minutes after the addition of Grb2. The first image in green is from EGFR^TAIL^. The second image in magenta is from Grb2. The third image is the merged image from EGFR^TAIL^ and Grb2.

The EGFR^TAIL^:Grb2 condensation is reversible and can be dispersed by addition of phosphatase. Addition of the phosphatase YopH (1 μM YopH and 6 μM Grb2 in bulk solution) rapidly disrupts the EGFR^TAIL^:Grb2 condensate, returning the system to a homogeneous and mobile distribution of EGFR^TAIL^ within 1 minute (see Figure 3). This confirms that even in the condensed state, sufficient dynamic unbinding and rebinding between Grb2 and the phosphorylated tyrosine sites on EGFR occurs to enable ready access by phosphatases (similar dynamic behavior has been observed in the LAT protein condensates (5)).

**Figure 3.**
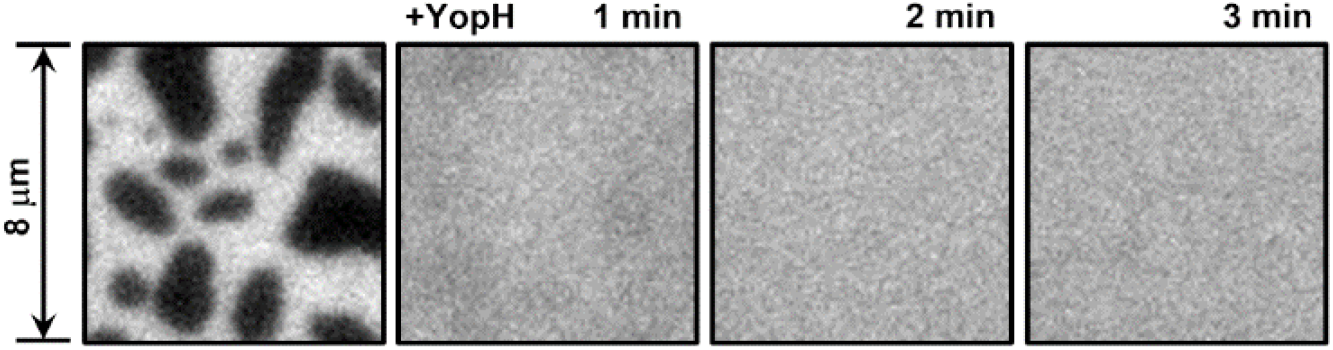
EGFR^TAIL^:Grb2 phase transition is reversed by phosphotase. The first image is the TIRF image of EGFR^TAIL^ condensation taken at 15 minutes after the addition of Grb2 to phosphorylated EGFR^TAIL^ on the supported bilayer. The second image was taken after the addition of 1 μM phosphatase, YopH. EGFR^TAIL^:Grb2 condensate was rapidly dissolved in 1 minute. The last image taken at 3 minutes after the addition of YopH shows uniformly distributed EGFR^TAIL^ (also see Figure S1).

Formation of the EGFR^TAIL^:Grb2 protein condensates requires a multivalent crosslinking interaction. In previous studies of the LAT:Grb2:SOS protein condensate, SOS was presumed to be the primary crosslinking element. In the experiments described here, with no SOS present, we hypothesized that crosslinking must be occurring through Grb2 directly. The crystal structure of Grb2 reveals a dimeric assembly (PDB 1GRI) and Grb2 dimerization plays a functional role in FGFR signaling (32). In the case of FGFR signaling, phosphorylation at Y160 on Grb2 breaks the dimer and enables downstream signaling. The Grb2 Y160 residue is located in the Grb2 dimer interface and a Y160E Grb2 mutant (referred to as Grb2^Y160E^) has been reported to block dimerization (33). To test if a Grb2 dimer interaction is responsible for the EGFR^TAIL^:Grb2 condensation, we characterize the condensation with the Grb2^Y160E^ as well as phosphorylated wildtype Grb2 (pGrb2). Experimentally, wildtype Grb2 was phosphorylated by Hck and further analysis by mass spectrometry confirmed phosphorylation at Y160 (see table 1 in SI). In both cases, EGFR^TAIL^:Grb2 condensation is fully disrupted (Figure 4, A, B, and C, and Figure S2). A Grb2 dimer interface serves as the crosslinking element in the EGFR^TAIL^:Grb2 condensate and this interaction can be negatively regulated by phosphorylation of Grb2 at Y160. The condensate formed by Grb2 and the EGFR^TAIL^ is analogous to that formed by Grb2 and LAT, and we speculate that the same Grb2-mediated effects are also present in LAT condensates. Experiments with LAT and Grb2 confirm that a LAT:Grb2 condensate also forms in a manner very similar to what we have observed with the EGFR^TAIL^, that is, Grb2 alone can drive the condensation, without the addition of SOS or other crosslinking elements (Figure 4D). Control experiments without ATP confirm that the EGFR^TAIL^:Grb2 condensate is dependent on EGFR phosphorylation (Figure 4E).

**Figure 4.**
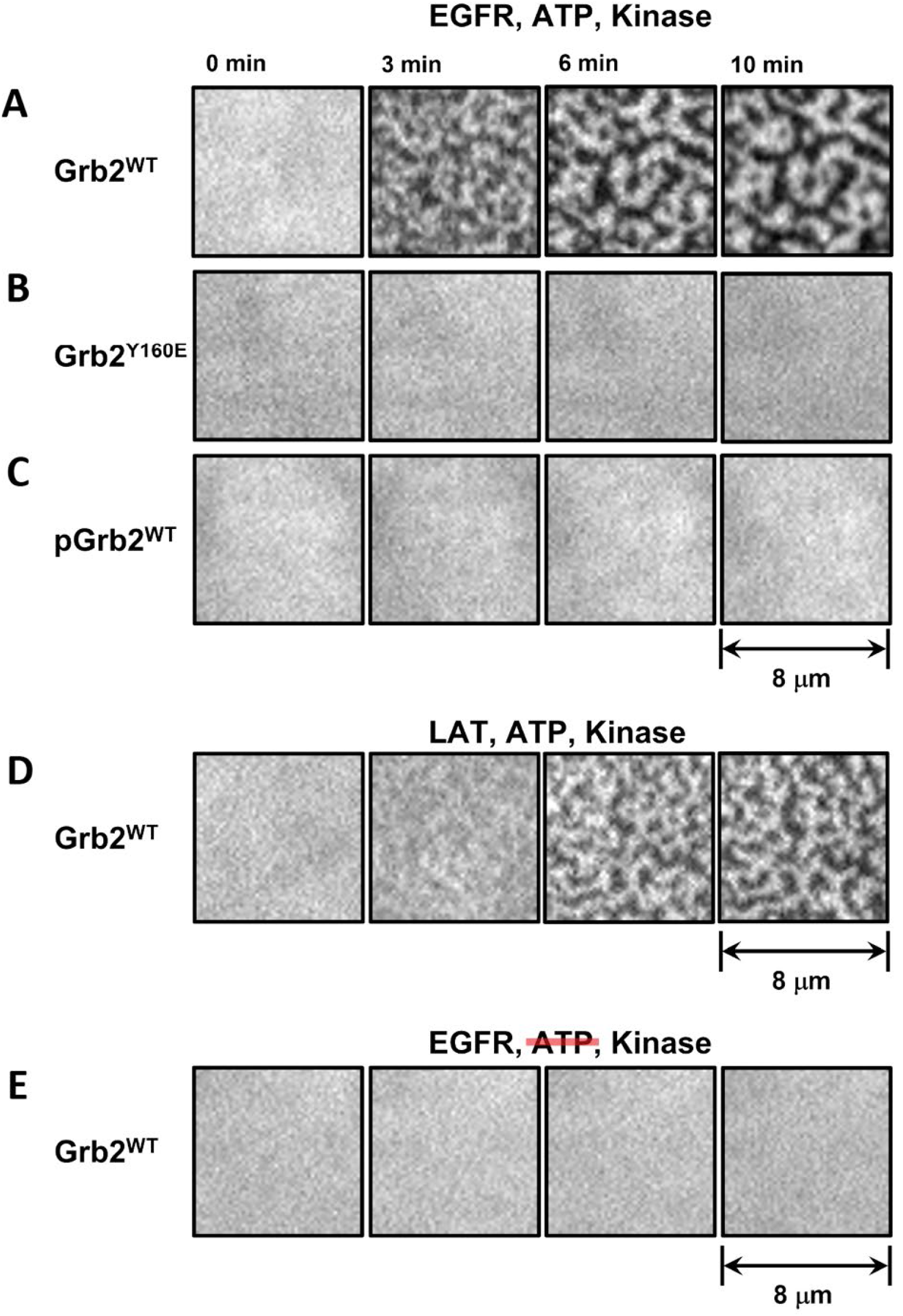
Grb2 dimerization as the key driving force to induce phase transition. TIRF images of EGFR showing the reconstitutions of (A) wildtype Grb2 and EGFR^TAIL^. Phase transition is observed at 3 minutes. (B) Grb2^Y160E^ and EGFR^TAIL^. EGFR^TAIL^ remains uniformly distributed. Phase transition is not detected. (C) pGrb2 and EGFR^TAIL^. Phase transition is not detected. (D) wildtype Grb2 and LAT. Phase transition of LAT:Grb2 is observed in 6 minutes. (E) wildtype Grb2 and EGFR^TAIL^ without ATP. Phase transition is not detected. In (A-E), Hck is the kinase used to phosphorylate EGFR^TAIL^.

### EGFR condensation enhances Ras activation by SOS

To measure the effects of EGFR condensation on downstream signaling, we reconstitute the signaling pathway from EGFR to Ras. This reconstitution is depicted schematically in Figure 5 and includes the EGFR^TAIL^, Grb2, phosphatidylinositol-4,5 bisphosphate (PIP_2_) (which participates in SOS autoinhibition release), full-length SOS (SOS^FL^), Ras, and the Ras binding domain of Raf (referred as RBD), which provides a final readout of Ras activation by its selective recruitment to RasGTP (10). Ras is anchored to the membrane via MCC-PE lipids using maleimide chemistry following well-established methods (43–45). Prior to any experiments, membrane-associated Ras is loaded with GDP to establish a starting condition of fully inactive Ras (see methods). In these experiments, Grb2 and the SOS^FL^ protein are maintained at sufficiently low concentrations that their presence alone is insufficient to trigger a condensation phase transition.

**Figure 5.**
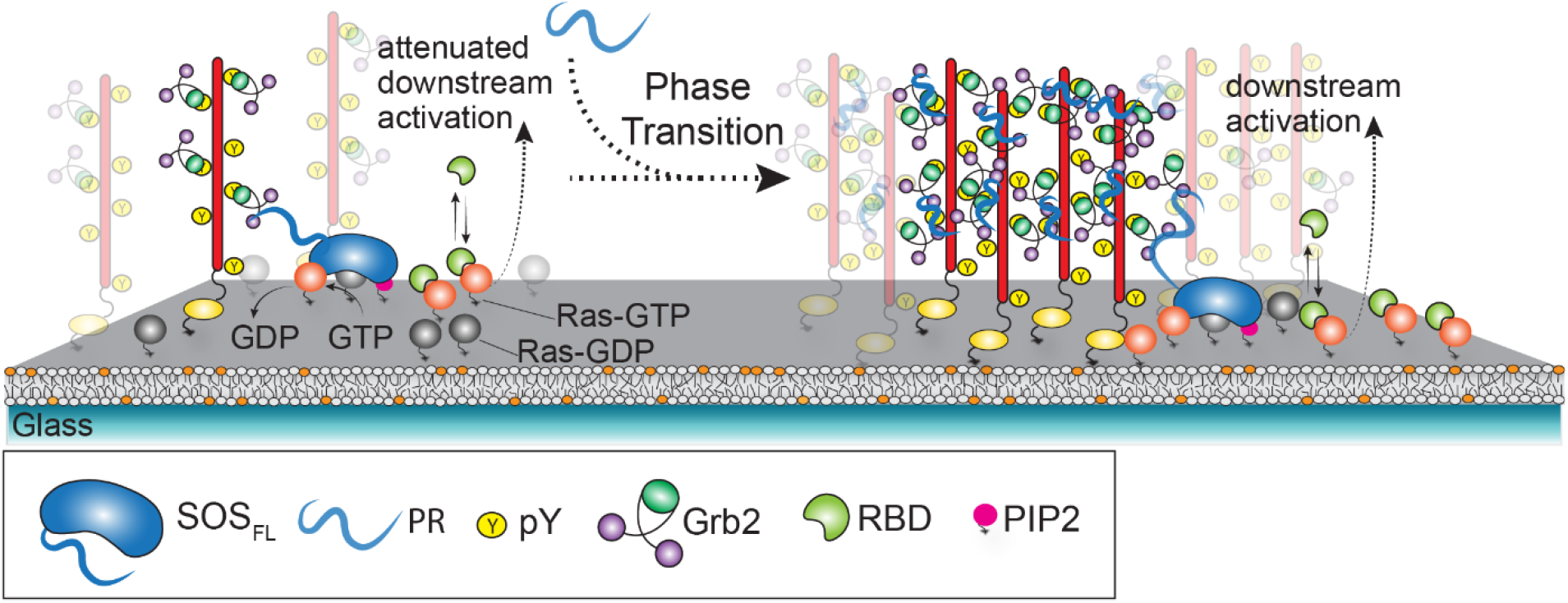
The schematic of downstream signaling modulated by SOS^Pr^ titrated EGFR^TAIL^:Grb2 phase transition. SOS^PR^ was used as the strong crosslinker to induce the phase transition of the molecule assembly, EGFR^TAIL^:Grb2:SOS, at different levels. To initiate the downstream signaling, Grb2, SOS^PR^, GTP (1 mM), Alexa Fluor 555 labeled SOS^FL^ and Alexa Fluor 647 labeled RBD (50 μM) were added together to the bilayer. Once SOS^FL^ was activated, the activated SOS^FL^ would stay on the membrane processively activating Ras. The activated Ras (Ras-GTP) is detected by RBD. The downstream signaling at the activation of Ras is read by the fluorescence intensity of RBD on the bilayer. Also see Ras activation in Materials and Methods.

To control the phase transition, we titrate in the proline-rich domain of SOS (residues 1051-1333; referred as SOS^PR^). SOS^PR^ is a strong crosslinker of Grb2 and has been previously used to control the LAT:Grb2:SOS condensation (10), and behaves similarly with the EGFR^TAIL^. Unlike SOS^FL^, SOS^PR^ lacks the catalytic domains and thus cannot contribute to Ras activation. To initiate the Ras activation reaction, SOS^FL^ (Alexa Fluor 555 labeled), Grb2, and SOS^PR^ are added in various ratios together with GTP (1 mM) and Alexa Fluor 647-labeled RBD (50 µM) to the pre-phosphorylated EGFR^TAIL^ and RasGDP on the supported membrane. Subsequently, Grb2 binds to the phosphotyrosine sites on the EGFR^TAIL^, recruiting both SOS^FL^ and SOS^PR^, and a lateral protein condensation phase transition proceeds to various degrees depending on the specific protein concentrations in each experiment. At the membrane, full-length SOS further engages PIP_2_ through its PH domain and Ras (GDP or GTP state) through its allosteric binding site (46, 47), ultimately releasing autoinhibition and beginning to catalyze the nucleotide exchange reaction of RasGDP to RasGTP. RasGTP levels are read out in real time through the selective binding of Alexa Fluor 647-labeled RBD to RasGTP. We use a modified RBD construct that exhibits fast and reversible binding kinetics (10), and achieves ∼5s time resolution for tracking RasGTP.

Ras activation trajectories are tracked as a function of EGFR^TAIL^ condensation state for two different overall SOS^FL^ concentrations (0.2 and 2 nM). Data plotted in Figures 6A and 6B illustrate the overall Ras activation trajectories as well as the calibrated, per-molecule SOS activities for each condition. Representative images of the corresponding EGFR^TAIL^ condensation state are illustrated in Figure 6C. For both SOS^FL^ concentrations, titrating in SOS^PR^ leads to an abrupt enhancement in Ras activation after macroscopic condensation is achieved, as seen by the sharp divergence of the fastest trajectory (red) from trajectories for the other conditions in each case. This enhancement of Ras activation in EGFR^TAIL^ condensates is observed in the calibrated, per-molecule, SOS activities (Figures 6A and 6B upper panels) as well as in overall reaction trajectories (lower panels). Thus, in addition to recruitment of more SOS to the condensates (e.g. through multivalent avidity effects), each SOS^FL^ molecule also activates more Ras over the same time period compared with uncondensed state of the EGFR^TAIL^. Also prominent in the fastest overall reaction trajectories from each SOS^FL^ concentration is RasGTP-driven positive feedback, which is evidenced by the positive curvature in the trajectories (20, 38, 48). In these reactions, a constant catalytic rate from SOS^FL^ will result in a Ras activation trajectory with strictly negative curvature as progressively less RasGDP is available to be activated.

**Figure 6.**
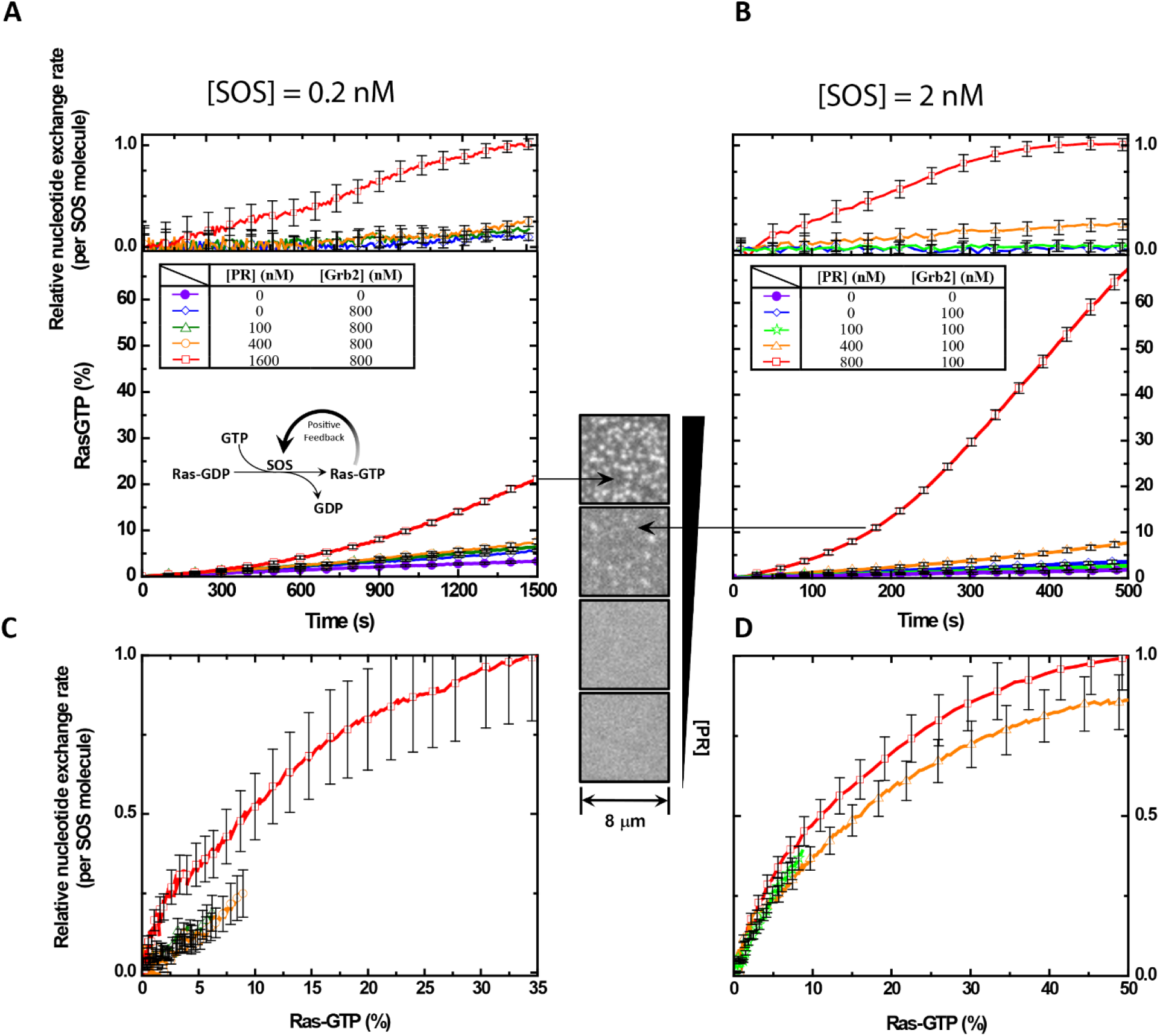
Ras activation modulated by EGFR^TAIL^:Grb2 phase transition. (A) The kinetic traces of Ras activation and the relative nucleotide exchange as the function at [SOS] = 0.2 nM. (B) The kinetic traces of Ras activation and the relative nucleotide exchange as the function of time at [SOS] = 2 nM. (C) The plot of the relative nucleotide exchange rate against the activated Ras showing the positive feedback of SOS at [SOS] = 0.2 nM. (D) The plot of the relative nucleotide exchange rate against the activated Ras showing the positive feedback of SOS at [SOS] = 2 nM.

Further insight into how the EGFR condensate may be facilitating SOS activation of Ras can be obtained by examining the reaction rate as a function of RasGTP density. Plots of the relative overall nucleotide exchange rate as a function of RasGTP density for each condition in Figures 6A and 6B are presented in Figures 6C and 6D. Strong rate enhancements with increasing RasGTP are evident in all cases, again revealing RasGTP-driven positive feedback. The EGFR condensate-driven enhancement in Ras activation, which is dominant in the time trajectories of these reactions, is much less obvious when rates are mapped to RasGTP levels. At the higher SOS^FL^ concentration (Figure 6E), the EGFR condensate effect is essentially undetectable, with all trajectories nearly overlaying each other. At the lower SOS^FL^ concentration (Figure 6D), a modest increase (roughly double) in reaction rate is evident for the condensed state (red) vs. the other conditions. In the related LAT condensate system, Ras activation by SOS has been shown to follow a first passage time mechanism, in which the overall system kinetics are predominantly set by the time it takes to get the first few SOS molecules activated (49) (although effects of LAT condensation were not addressed in that study). Ras activation measurements presented here suggest EGFR condensation is impacting the initiation of Ras activation more so than it is modulating steady-state aspects of the reaction. More efficient activation of the first few SOS molecules allows the RasGTP-driven positive feedback to kick in earlier, leading to a substantial difference in overall Ras activation.

## Discussion and Conclusion

The protein condensate of the EGFR^TAIL^ described here exhibits striking similarities with the LAT protein condensate, with respect to its phase transition properties as well as the way it impacts Ras activation. However, the context of the LAT protein condensation in response to TCR activation differs dramatically from EGFR signaling with respect to the copy numbers of molecules involved. T cells have near single molecule antigen sensing capabilities (50–53), and can activate with only few triggered TCRs. In contrast, EGFR stimulation often involves thousands or even millions of activated receptors per cell (54–56). Among molecules in the EGFR-MAPK pathway, SOS is expressed at very low levels and is likely a limiting component (56). Thus unlike the LAT:Grb2:SOS condensate, an EGFR:Grb2:SOS condensate would seem implausible if SOS was a key crosslinking element that needed to be present in stoichiometric ratios with EGFR. Observations of a two-component EGFR:Grb2 condensate that we report here, with Grb2 dimerization serving as the primary crosslinker, relieve SOS of its requisite crosslinking role. We are then free to envision physiological EGFR condensates, primarily scaffolded by Grb2 dimer interactions, which contain substantially sub-stochiometric copy numbers of SOS relative to the numbers of EGFR molecules. EGFR condensation nonetheless exhibits significant control over the ability of SOS to activate Ras.

More broadly, the EGFR condensate establishes a layer of the EGFR signal transduction mechanism, proximal to ligand-mediated EGFR activation itself, that offers regulatory control over the efficiency of downstream signal propagation. EGFR condensation may be influenced by other physical and chemical parameters, such as local mobility of EGFR, membrane topographical features, or other molecules participating in the condensation (57–59). Additionally, the fact that Grb2 dimerization is disrupted by Y160 phosphorylation reveals a potential active regulatory mechanism by which EGFR condensation could be curtailed. Since Grb2 in condensates is in constant exchange with cytosolic Grb2 (5), such a mechanism would require a cell-wide shift in Grb2 phosphorylation levels; it remains an open question if such regulation is utilized under physiological conditions. An intriguing side effect of Grb2 phosphorylation as a potential negative regulator is that tyrosine kinase inhibitors, such as are widely used as cancer therapeutics in EGFR-driven cancers (60), could also inhibit this negative regulatory mechanism—conceivably interfering with their desired effects.

## Materials and Methods

### Chemicals

1,2-dioleoyl-sn-glycero-3-phosphocholine (DOPC) and 1,2-dioleoyl-sn-glycero-3-[(N-(5-amino-1-carboxypentyl)iminodiacetic acid)succinyl] (nickel salt) (Ni^2+^-NTADOGS) were purchased from Avanti Polar Lipids. Alexa Fluor 488, Alexa Fluor 555, and Alexa Fluor 647 maleimide dyes were purchased from ThermoFisher. Bovine Serum Albumin (BSA), (±)-6-Hydroxy-2,5,7,8-tetramethylchromane-2-carboxylic acid (Trolox), Catalase, 2-Mercaptoethanol (BME), NiCl_2_, H_2_SO_4_ and ATP were purchased from Sigma-Aldrich. Glucose Oxidase was purchased from Serva. Tris(2-carboxyethyl)phosphine (TCEP) was brought from Thermo Scientific. Glucose and H_2_O_2_ were from Fisher Scientific. MgCl_2_ was from EMD Chemicals. Tris buffer saline (TBS) was purchased from Corning.

### Protein purification

#### EGFR^TAIL^

C-terminal region of EGFR^TAIL^ (residues 1015-1210, where the signal peptide of EGFR is included in numbering) fused to a N-terminal His_6_ and a SUMO tag with a TEV cleaving site was cloned into pProEX-HTb. The plasmid was transformed into BL21(DE3) *Escherichia coli*. The transformed cells were grown at 37°C in 1L Terrific Broth media until OD_600_ reached 0.7. The culture was induced with 1 mM isopropyl β-D-1-thiogalactopyranoside (IPTG) and let grow overnight at 18 °C. After overnight expression, the culture is spun down for 15 minutes at 4000 rpm and resuspended in 15-30 mL of NiNTA buffer (500 mM NaCl, 20 mM Tris-HCl pH 8.5, 20 mM Imidazole). The EGFR^TAIL^ expressing cells were lysed by sonication. The insoluble fraction of the lysate was separated by ultracentrifugation. The supernatant was applied to a HisTrap FF (GE) column. EGFR^TAIL^ was eluted by eluting buffer (500 mM NaCl, 20 mM Tris-HCl pH 8.5, 500 mM Imidazole). The elution fractions was concentrated by an Amicon Ultra centrifugal filter unit (30k Da MWCO, Millipore). The concentrated Grb2 is loaded onto an S75 30/300 (GE Healthcare) column equilibrated in Buffer A (150 mM NaCl, 20 mM Tris-HCl pH 8.0, 1 mM TCEP). Finally, purified EGFR^TAIL^ was then aliquoted and stored at -80 °C after flash freezing for further labeling reactions.

#### Grb2

pET-28 plasmid containing full-length Grb2 fused to N-terminal His6 was transformed into BL21(DE3) *Escherichia coli*. The transformed cells were grown at 37°C in 1L Terrific Broth media until OD_600_ reached 0.7. The culture was induced with 1 mM isopropyl β-D-1-thiogalactopyranoside (IPTG) and let grow overnight at 18 °C. After overnight expression, the culture is spun down for 15 minutes at 4000 rpm and resuspended in 15-30 mL of NiNTA buffer (500 mM NaCl, 20 mM Tris-HCl pH 8.5, 20 mM Imidazole, 5% Glycerol). The Grb2 expressing cells were lysed by sonication with addition of 1mM PMSF and 0.1 mM BME. The insoluble fraction of the lysate was spun down at 16500 rpm for 45 minutes at 4 °C. The supernatant was applied to a HisTrap FF (GE) column. Grb2 was eluted by eluting buffer (500 mM NaCl, 20 mM Tris-HCl pH 8.5, 500 mM Imidazole, 5% Glycerol). The elution fraction was loaded onto a Desalting column equilibrated in Buffer A (150 mM NaCl, 20 mM Tris-HCl pH 8.0, 1 mM TCEP 5% Glycerol). The fraction of the Grb2 peak was collected. Grb2 was incubated with TEV protease overnight to cleave the His tag off. The cleaved Grb2 was reapplied to a HisTrap column. The flow-through was collected and concentrated by an Amicon Ultra centrifugal filter unit (10k Da MWCO). The concentrated Grb2 is loaded onto an S75 30/300 (GE Healthcare) column equilibrated in Buffer A. The fractions of Grb2 monomer were collected and concentrated. Grb2 was further equilibrated at 37 °C for 10 min to reestablish the equilibrium between the monomer and dimer. Finally, Grb2 was then aliquoted and stored at -80 °C after flash freezing.

#### Ras

H-Ras S118C containing residue 1-181 fused to N-terminal His_6_ was purified. The N-terminal His_6_ tag was removed by TEV protease. The procedures were described in the previous work (61).

#### SOS^FL^

Full-length SOS protein was prepared via a split intein approach. The N- and the C-terminal fragments of human SOS1 were expressed in BL21 (DE3) bacteria and purified separately. The two fragments with high purity were ligated by the intein reaction to generate SOS^FL^. The details of the design for the intein reaction, expression and purification are described in the previous work. (10).

#### SOS^PR^

pETM vector containing the ORF, His6-MBP-(Asn)10-TEV-SOS1 (residue 1051–1333, human) proline-rich domain was transformed into BL21 (DE3) bacteria. The expression and purification can be found in the previous work (5).

#### RBD-K65E

pETM11 vector containing the ORF, his_6_-GST-PreScission-SNAPtag-Raf1 RBD (residue 56-131, K65E) derived from the Raf-1 human gene was transformed into BL21 (DE3) bacteria. RBD was expressed and purified as described previously (10).

### Protein labeling

Proteins were first diluted to 100 µM. Alexa Fluor 488, 555 or 647 maleimide dye was dissolved in anhydrous DMSO to make a working stock that is 10-20mM. The dye was added to the protein within 3 fold molar excess over protein concentration to avoid overlabeling. The reaction mixture is incubated at the room temperature for 1 hr. To quench the labeling reaction, Dithiothreitol (DTT) was added to the reaction mixture at the final concentration of 10 mM. Amicon Ultra centrifugal filter unit was used to wash the free dye away by adding more buffer while concentrating the protein. The concentrated protein was loaded to an analytical grade size exclusion column (S75, GE Healthcare) for the further purification. The fractions of the purified protein were collected and concentrated. Finally, the labeled protein was aliquoted and stored at -80 °C after flash freezing.

### Supported lipid bilayers (SLB)

SLBs were formed by rupturing small unilamellar vesicles (SUVs). SUVs were made by mixing DOPC, Ni^2+^-NTA-DOGS, PE MCC, and PIP2 lipids (92:4:2:2 by molar percent) in chloroform. Chloroform of solution was evaporated using a rotary evaporator for 10 min at 40°C to obtain the dried lipid films which were further dried by purging N_2_ for 10 min. The vesicle solution was formed from the hydration of the dried lipids by H_2_O with the concentration of 1 mg/ml using vortexing. SUVs were made by sonicating the vesicle solution for 90 s in an ice-water bath. SLBs were prepared on the glass substrate cleaned by piranha etch (Ibidi glass coverslips, bottom thickness 170 µm +/– 5 µm) combined with the flow chamber (sticky-Slide VI 0.4, Ibidi, Planegg, Germany). The SUVs were mixed with 10 mM TBS buffer at pH 7.4 (1:1 ratio by volume) and injected into the chamber for 30 min of incubation to make SLBs. The incubation of 1 mg/mL BSA in TBS buffer for 10 min was used to block the defects in SLBs. H-Ras was anchored on PE MCC lipids through the maleimide chemistry by incubating 0.5 mg/ml H-Ras with SLBs for 3 hr. The free PE MCC lipids were quenched by 5 mM BME. EGFR^TAIL^ and Hck were anchored on SLBs via his-tag – Ni^2+^NTA chemistry by incubating EGFR^TAIL^ and Hck at 150 and 10 nM for 30 min. During this step, 1 mM GDP was included to ensure that Ras is loaded with GDP. To phosphorylate the tyrosine sites of EGFR^TAIL^, 1 mM ATP and 5 mM MgCl_2_ in TBS were added to SLBs and incubated for 10 min. The mobility of the membrane-bound proteins was examined by fluorescent recovery after photobleaching (FRAP). The densities of the membrane proteins were determined by the averaged intensities of the epifluorescence images via the calibration curve between the intensity of the image and the density of the protein measured by the fluorescence correlation spectroscopy (FCS) described previously (43).

### Total internal reflection fluorescence (TIRF) microscopy

An Eclipse Ti inverted microscope (Nikon) with a TIRF system and iXon EMCCD (Electron Multiplying Charged-Coupled Device) camera (Andor Technology) was used for imaging. TIRF microscopy was performed with a Nikon 100 × 1.49 numerical aperture oil-immersion TIRF objective, a TIRF illuminator, a Perfect Focus system, a motorized stage and a U-N4S 4-laser unit as a laser source (Nikon). The laser unit has the solid state lasers for the 488 nm, 561 nm and 640 nm channels, and was controlled using a built-in acousto-optic tunable filter (AOTF). The laser powers of 488 nm laser, 561 nm laser, and 640 nm laser were set to 5.2, 6.9, and 7.8mW measured with the field aperture fully opened. The 405/488/561/638 nm Quad TIRF filter set (Chroma Technology Corp., Rockingham, Vermont) along with supplementary emission filters of 525/50 m, 600/50 m, 700/75 m for 488 nm, 561 nm, 640 nm channel was used respectively. Images are acquired using the Nikon NIS-Elements software with the exposure time of 50 milliseconds.

### Ras Activation and Imaging Analysis

SOS^PR^ was used as the strong crosslinker to induce the phase transition of the molecule assembly, EGFR^TAIL^:Grb2:SOS, at different levels while the concentrations of Grb2 were maintained constant avoiding the bias in signal outputs due to the competition for Grb2. SOS^PR^ without the catalytic domain for the nucleotide exchange of Ras would not contribute to the Ras activation done by SOS^FL^. To initiate the downstream signaling, Grb2, SOS^PR^, GTP (1 mM), Alexa Fluor 555 labeled SOS^FL^ and Alexa Fluor 647 labeled RBD (50 μM) were added together to the bilayer. While the phase transition of EGFR^TAIL^ was occurring, SOS was recruited by Grb2. Once SOS^FL^ was activated, the activated SOS^FL^ would stay on the membrane and start to processively activate Ras. The activated Ras (Ras-GTP) on the membrane was further detected by RBD in the bulk solution via the interaction between Raf and activated Ras. The downstream signaling at the activation of Ras was read by the fluorescence intensity of RBD on the bilayer using TIRF images. At the end of each Ras activation trace, 20 nM SOS^cat^, the catalytic domain of SOS^FL^, was used to fully activate Ras on SLB. The averaged intensity of the RBD TIRF image corresponding to total Ras molecules on SLB was used as the normalization factor to calibrate different Ras densities over different traces. SOS on the membrane can be tracked at the same time while the fluorescence intensity of RBD on the bilayer was being monitored. The Ras activation by SOS was allowed to interact directly with the phase transition of EGFR^TAIL^. The concentrations of Grb2 (800 nM and 100 nM) were chosen to modulate the recruitment of SOS by Grb2 making the time scale of Ras activation comparable within 30 min between the two data sets of low and high [SOS].

Single-molecule images of SOS^FL^ were tracked by an ImageJ plugin, TrackMate (62), to obtain the number of activated SOS^FL^ on the membrane. SOS^FL^ molecules were localized with the detector of Difference of Gaussian (DoG). The initial diameter was set to six pixels. Typically, 50–3000 molecules were detected per image. In this study, the Ras activation modulated by EGFR^TAIL^ phase transition at both low and high [SOS] (0.2 and 2 nM) were interrogated separately. Dividing the instantaneous increase of activated Ras by the number of SOS on the bilayer gave the relative nucleotide exchange rate of SOS as a function of time. To study the positive feedback of SOS^FL^, the relative nucleotide exchange rates of SOS^FL^ were plotted against activated Ras (Ras-GTP %).

## Supporting Information

**Figure S1.**
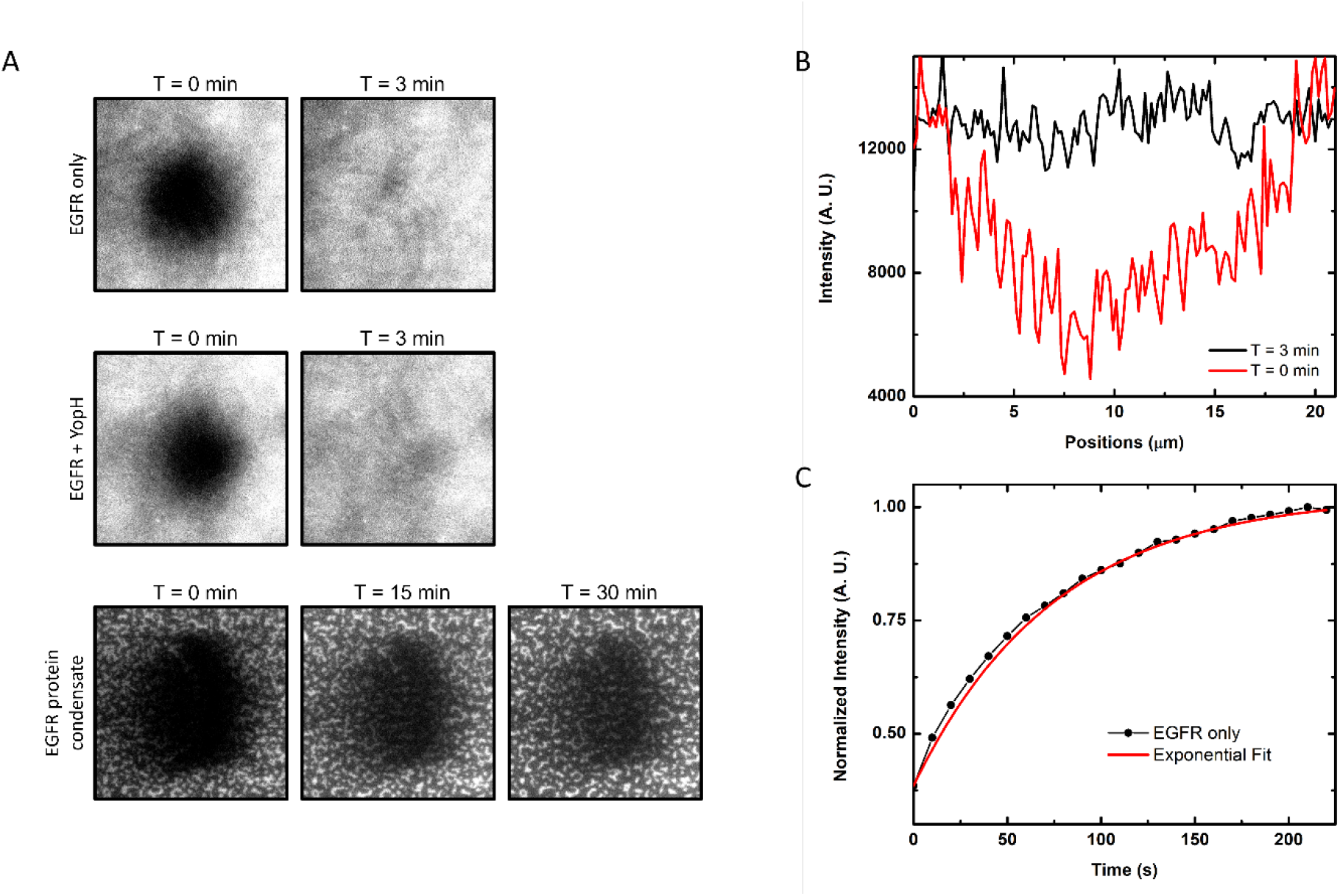
FRAP experiments of EGFR. (A) FRAP images. Top: EGFR on the bilayer, middle: EGFR^TAIL^ after YopH treatment, bottom: EGFR^TAIL^:Grb2 protein condensate. (B) Intensity profile of FRAP image from EGFR^TAIL^ only at 0 and 3 min. (C) The recovery trace of EGFR^TIAL^ only. The red solid line is the exponential fit where τ_D_ = 73 s.

**Figure S2.**
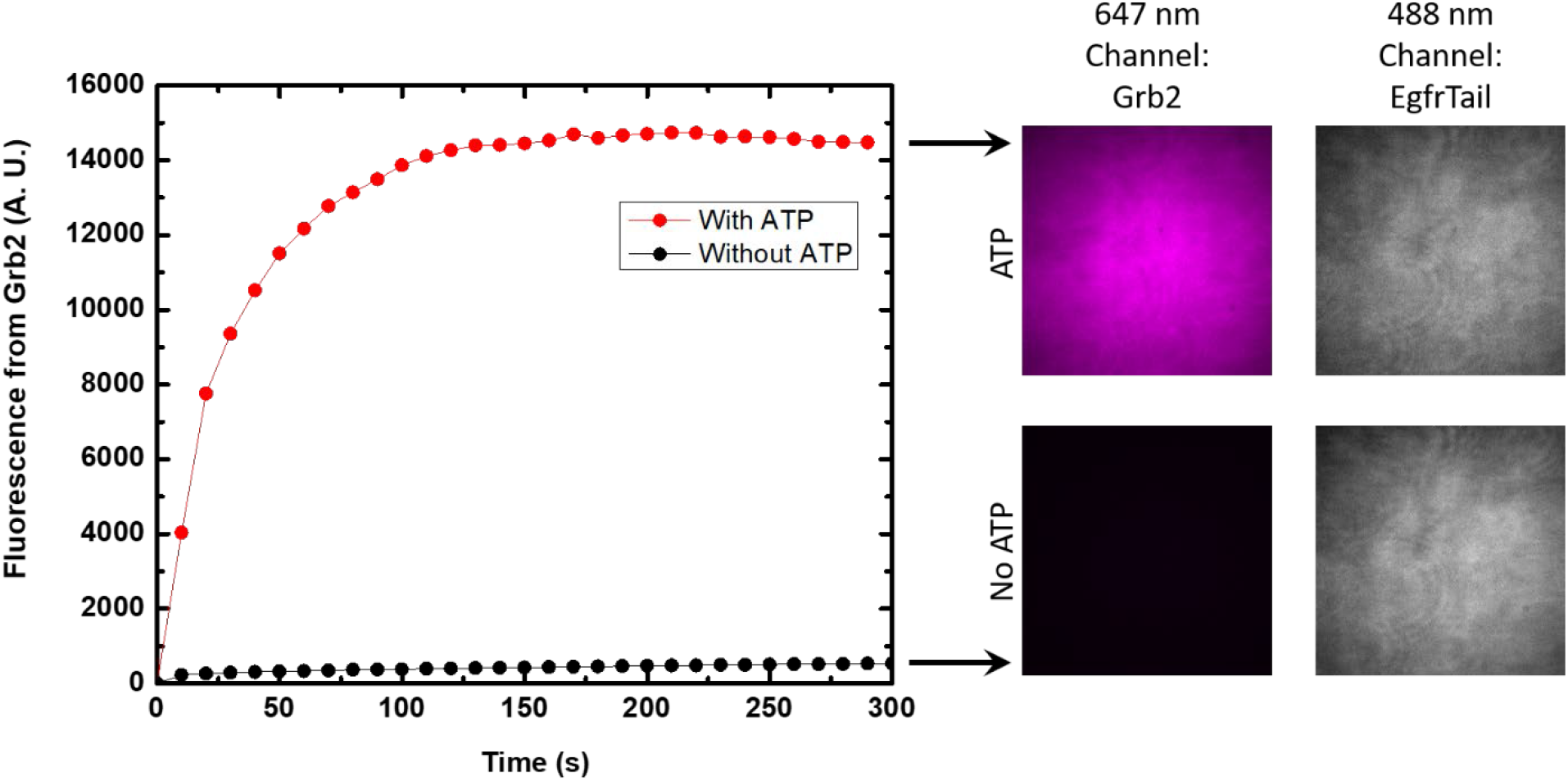
Grb2^Y160E^ responses to the phosphorylation of EGFR^TAIL^. Alexa Fluor 647 labeled Grb2^Y160E^ (6 μM, labeling efficiency of 48%) is added to phosphorylated EGFR^TAIL^ on the bilayer. The averaged fluorescence intensity from the TIRF images of Grb2^Y160E^ on the supported bilayer increases rapidly corresponding to the recruitment by EGFR^TAIL^ (the red curve in the left graph). The control experiment is shown by the black curve where EGFR^TAIL^ is not phosphorylated. The TIRF images from Grb2^Y160E^ (in magenta) and EGFR^TAIL^ (in gray) taken 10 minutes after the addition are shown at the right side of the figure. No phase transition is observed using Grb2^Y160E^.

**Table S1.**
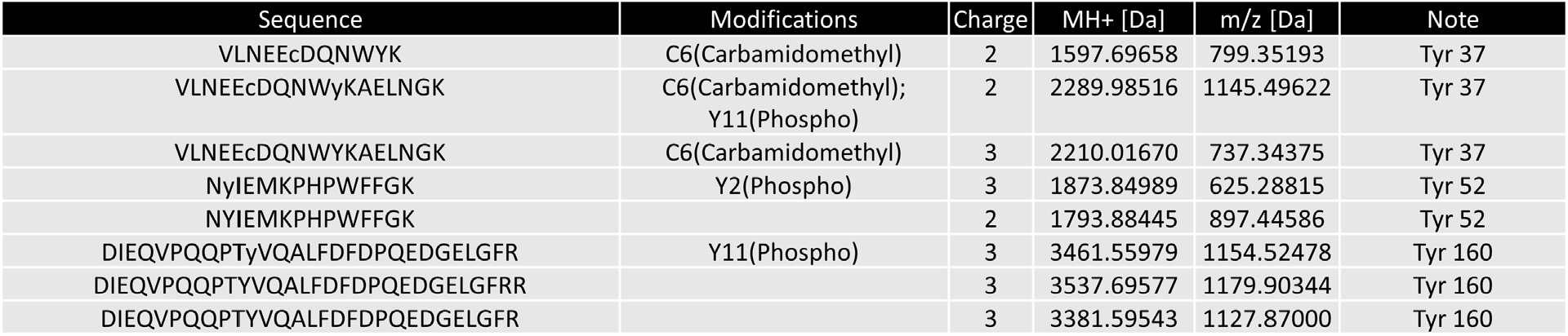
Three tyrosines of Grb2 show the phosphorylation by Hck. Grb2 has seven tyrosines. The peptide analysis of mass spectrum covers 85% of Grb2 sequence which includes six tyrosines in Grb2. The phosphorylated Grb2 sample was digested by trypsin overnight before the analysis by the mass spectrometer.

**Figure S3.**
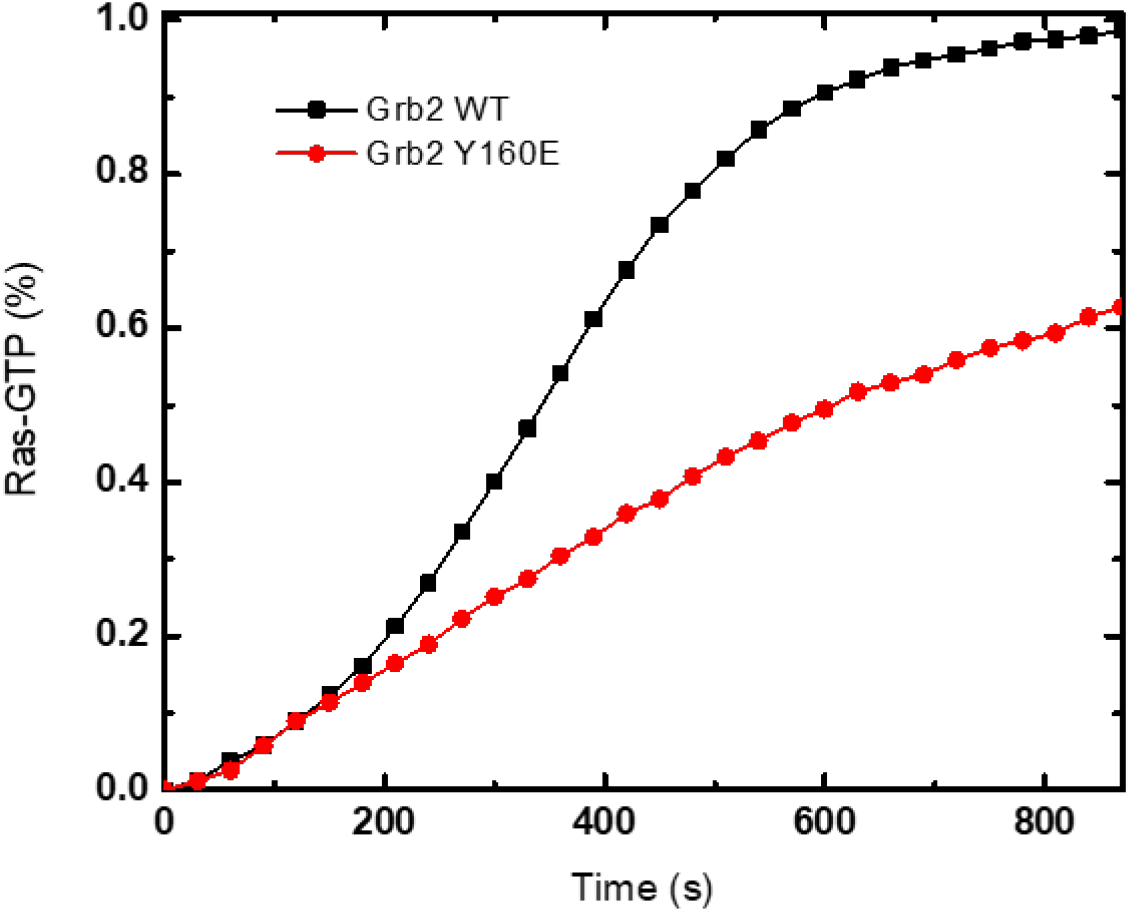
Ras activation between wildtype Grb2 and Grb2^Y160E^. In this Ras activation assay, the crosslinker, SOS^PR^ is not used. To initiate the downstream signaling, Grb2 (200 nM), GTP (1 mM), Alexa Fluor 555 labeled SOS^FL^ (4 nM) and Alexa Fluor 647 labeled RBD (50 μM) are added together to phosphorylated EGFR^TAIL^ on the supported bilayer. The activated Ras (Ras-GTP) is detected by RBD. Also see Ras activation in Materials and Methods.

## Bibliography

1. Y. Shin, C. P. Brangwynne, Liquid phase condensation in cell physiology and disease. Science 357, eaaf4382 (2017).

2. D. Hnisz, K. Shrinivas, R. A. Young, A. K. Chakraborty, P. A. Sharp, A phase separation model for transcriptional control. Cell 169, 13–23 (2017).

3. R. L. Kortum, et al., The ability of Sos1 to oligomerize the adaptor protein LAT is separable from its guanine nucleotide exchange activity in vivo. Sci. Signal. 6, ra99 (2013).

4. X. Su, et al., Phase separation of signaling molecules promotes T cell receptor signal transduction. Science 352, 595–599 (2016).

5. W. Y. C. Huang, et al., Phosphotyrosine-mediated LAT assembly on membranes drives kinetic bifurcation in recruitment dynamics of the Ras activator SOS. Proc Natl Acad Sci USA 113, 8218–8223 (2016).

6. L. Balagopalan, R. L. Kortum, N. P. Coussens, V. A. Barr, L. E. Samelson, The linker for activation of T cells (LAT) signaling hub: from signaling complexes to microclusters. J. Biol. Chem. 290, 26422–26429 (2015).

7. E. Sherman, V. Barr, L. E. Samelson, Super-resolution characterization of TCR-dependent signaling clusters. Immunol. Rev. 251, 21–35 (2013).

8. S. Banjade, M. K. Rosen, Phase transitions of multivalent proteins can promote clustering of membrane receptors. eLife 3, e04123 (2014).

9. L. B. Case, X. Zhang, J. A. Ditlev, M. K. Rosen, Stoichiometry controls activity of phase-separated clusters of actin signaling proteins. Science 363, 1093–1097 (2019).

10. W. Y. C. Huang, et al., A molecular assembly phase transition and kinetic proofreading modulate Ras activation by SOS. Science 363, 1098–1103 (2019).

11. Y. Zhang, T. G. Kutateladze, Liquid-liquid phase separation is an intrinsic physicochemical property of chromatin. Nat. Struct. Mol. Biol. 26, 1085–1086 (2019).

12. O. Beutel, R. Maraspini, K. Pombo-García, C. Martin-Lemaitre, A. Honigmann, Phase separation of zonula occludens proteins drives formation of tight junctions. Cell 179, 923–936. (2019).

13. Y. Xie, et al., Polarisome scaffolder Spa2-mediated macromolecular condensation of Aip5 for actin polymerization. Nat. Commun. 10, 5078 (2019).

14. N. F. Endres, K. Engel, R. Das, E. Kovacs, J. Kuriyan, Regulation of the catalytic activity of the EGF receptor. Curr. Opin. Struct. Biol. 21, 777–784 (2011).

15. R. Avraham, Y. Yarden, Feedback regulation of EGFR signalling: decision making by early and delayed loops. Nat. Rev. Mol. Cell Biol. 12, 104–117 (2011).

16. M. A. Lemmon, J. Schlessinger, K. M. Ferguson, The EGFR family: not so prototypical receptor tyrosine kinases. Cold Spring Harb. Perspect. Biol. 6, a020768 (2014).

17. E. Kovacs, J. A. Zorn, Y. Huang, T. Barros, J. Kuriyan, A structural perspective on the regulation of the epidermal growth factor receptor. Annu. Rev. Biochem. 84, 739–764 (2015).

18. Z. Wang, Erbb receptors and cancer. Methods Mol. Biol. 1652, 3–35 (2017).

19. A. J. Cantor, N. H. Shah, J. Kuriyan, Deep mutational analysis reveals functional trade-offs in the sequences of EGFR autophosphorylation sites. Proc Natl Acad Sci USA 115, E7303–E7312 (2018).

20. S. Boykevisch, et al., Regulation of ras signaling dynamics by Sos-mediated positive feedback. Curr. Biol. 16, 2173–2179 (2006).

21. I. Chung, et al., Spatial control of EGF receptor activation by reversible dimerization on living cells. Nature 464, 783–787 (2010).

22. P. Nagy, J. Claus, T. M. Jovin, D. J. Arndt-Jovin, Distribution of resting and ligand-bound ErbB1 and ErbB2 receptor tyrosine kinases in living cells using number and brightness analysis. Proc Natl Acad Sci USA 107, 16524–16529 (2010).

23. S. Saffarian, Y. Li, E. L. Elson, L. J. Pike, Oligomerization of the EGF receptor investigated by live cell fluorescence intensity distribution analysis. Biophys. J. 93, 1021–1031 (2007).

24. Y. Huang, et al., Molecular basis for multimerization in the activation of the epidermal growth factor receptor. eLife 5, e14107 (2016).

25. S. Sigismund, D. Avanzato, L. Lanzetti, Emerging functions of the EGFR in cancer. Mol. Oncol. 12, 3–20 (2018).

26. D. Westover, J. Zugazagoitia, B. C. Cho, C. M. Lovly, L. Paz-Ares, Mechanisms of acquired resistance to first-and second-generation EGFR tyrosine kinase inhibitors. Ann. Oncol. 29, i10–i19 (2018).

27. F. Kai, A. P. Drain, V. M. Weaver, The extracellular matrix modulates the metastatic journey. Dev. Cell 49, 332–346 (2019).

28. R. Thomas, Z. Weihua, Rethink of EGFR in cancer with its kinase independent function on board. Front. Oncol. 9, 800 (2019).

29. L. Zeng, I. Palaia, A. Šarić, X. Su, PLCγ1 promotes phase separation of T cell signaling components. J. Cell Biol. 220, e202009154 (2021).

30. W. Y. C. Huang, H.-K. Chiang, J. T. Groves, Dynamic Scaling Analysis of Molecular Motion within the LAT:Grb2:SOS Protein Network on Membranes. Biophys. J. 113, 1807–1813 (2017).

31. S. Sun, T. GrandPre, D. T. Limmer, J. T. Groves, Kinetic frustration by limited bond availability controls the LAT protein condensation phase transition on membranes. BioRxiv (2021) https://doi.org/10.1101/2021.12.05.471009.

32. C.-C. Lin, et al., Inhibition of basal FGF receptor signaling by dimeric Grb2. Cell 149, 1514–1524 (2012).

33. Z. Ahmed, et al., Grb2 monomer-dimer equilibrium determines normal versus oncogenic function. Nat. Commun. 6, 7354 (2015).

34. N. W. Gale, S. Kaplan, E. J. Lowenstein, J. Schlessinger, D. Bar-Sagi, Grb2 mediates the EGF-dependent activation of guanine nucleotide exchange on Ras. Nature 363, 88–92 (1993).

35. H. Sondermann, et al., Structural analysis of autoinhibition in the Ras activator Son of sevenless. Cell 119, 393–405 (2004).

36. M. A. Lemmon, J. Schlessinger, Cell signaling by receptor tyrosine kinases. Cell 141, 1117–1134 (2010).

37. P. Bandaru, Y. Kondo, J. Kuriyan, The Interdependent Activation of Son-of-Sevenless and Ras. Cold Spring Harb. Perspect. Med. 9, a031534 (2019).

38. L. Iversen, et al., Molecular kinetics. Ras activation by SOS: allosteric regulation by altered fluctuation dynamics. Science 345, 50–54 (2014).

39. S. M. Christensen, et al., One-way membrane trafficking of SOS in receptor-triggered Ras activation. Nat. Struct. Mol. Biol. 23, 838–846 (2016).

40. Y. K. Lee, et al., Mechanism of SOS PR-domain autoinhibition revealed by single-molecule assays on native protein from lysate. Nat. Commun. 8, 15061 (2017).

41. J. K. Chung, et al., K-Ras4B Remains Monomeric on Membranes over a Wide Range of Surface Densities and Lipid Compositions. Biophys. J. 114, 137–145 (2018).

42. W. Y. C. Huang, J. A. Ditlev, H.-K. Chiang, M. K. Rosen, J. T. Groves, Allosteric modulation of grb2 recruitment to the intrinsically disordered scaffold protein, LAT, by remote site phosphorylation. J. Am. Chem. Soc. 139, 18009–18015 (2017).

43. W.-C. Lin, et al., H-Ras forms dimers on membrane surfaces via a protein-protein interface. Proc Natl Acad Sci USA 111, 2996–3001 (2014).

44. J. K. Chung, Y. K. Lee, H. Y. M. Lam, J. T. Groves, Covalent Ras Dimerization on Membrane Surfaces through Photosensitized Oxidation. J. Am. Chem. Soc. 138, 1800–1803 (2016).

45. S. M. Christensen, et al., Monitoring the Waiting Time Sequence of Single Ras GTPase Activation Events Using Liposome Functionalized Zero-Mode Waveguides. Nano Lett. 16, 2890–2895 (2016).

46. C. Zhao, G. Du, K. Skowronek, M. A. Frohman, D. Bar-Sagi, Phospholipase D2-generated phosphatidic acid couples EGFR stimulation to Ras activation by Sos. Nat. Cell Biol. 9, 706–712 (2007).

47. S. M. Margarit, et al., Structural evidence for feedback activation by Ras.GTP of the Ras-specific nucleotide exchange factor SOS. Cell 112, 685–695 (2003).

48. A. A. Lee, et al., Stochasticity and positive feedback enable enzyme kinetics at the membrane to sense reaction size. Proc Natl Acad Sci USA 118, e2103626118 (2021).

49. W. Y. C. Huang, S. Alvarez, Y. Kondo, J. Kuriyan, J. T. Groves, Relating cellular signaling timescales to single-molecule kinetics: A first-passage time analysis of Ras activation by SOS. Proc Natl Acad Sci USA 118, e2103598118 (2021).

50. Y. Sykulev, M. Joo, I. Vturina, T. J. Tsomides, H. N. Eisen, Evidence that a single peptide-MHC complex on a target cell can elicit a cytolytic T cell response. Immunity 4, 565–571 (1996).

51. D. J. Irvine, M. A. Purbhoo, M. Krogsgaard, M. M. Davis, Direct observation of ligand recognition by T cells. Nature 419, 845–849 (2002).

52. G. P. O’Donoghue, R. M. Pielak, A. A. Smoligovets, J. J. Lin, J. T. Groves, Direct single molecule measurement of TCR triggering by agonist pMHC in living primary T cells. eLife 2, e00778 (2013).

53. J. J. Y. Lin, et al., Mapping the stochastic sequence of individual ligand-receptor binding events to cellular activation: T cells act on the rare events. Sci. Signal. 12, eaat8715 (2019).

54. F. Zhang, et al., Quantification of epidermal growth factor receptor expression level and binding kinetics on cell surfaces by surface plasmon resonance imaging. Anal. Chem. 87, 9960–9965 (2015).

55. N. F. Endres, et al., Conformational coupling across the plasma membrane in activation of the EGF receptor. Cell 152, 543–556 (2013).

56. T. Shi, et al., Conservation of protein abundance patterns reveals the regulatory architecture of the EGFR-MAPK pathway. Sci. Signal. 9, rs6 (2016).

57. R. Parthasarathy, C. Yu, J. T. Groves, Curvature-modulated phase separation in lipid bilayer membranes. Langmuir 22, 5095–5099 (2006).

58. Y. Kaizuka, J. T. Groves, Bending-mediated superstructural organizations in phase-separated lipid membranes. New J. Phys. 12, 095001 (2010).

59. J. K. Chung, et al., Coupled membrane lipid miscibility and phosphotyrosine-driven protein condensation phase transitions. Biophys. J. 120, 1257–1265 (2021).

60. A. Ayati, et al., A review on progression of epidermal growth factor receptor (EGFR) inhibitors as an efficient approach in cancer targeted therapy. Bioorg. Chem. 99, 103811 (2020).

61. J. Gureasko, et al., Membrane-dependent signal integration by the Ras activator Son of sevenless. Nat. Struct. Mol. Biol. 15, 452–461 (2008).

62. J.-Y. Tinevez, et al., TrackMate: An open and extensible platform for single-particle tracking. Methods 115, 80–90 (2017).

